# A Generalizable Machine Learning Framework for cfDNA based Early Detection of Hepatocellular Carcinoma: a Feasibility Study with Preclinical Validation

**DOI:** 10.1101/2025.06.09.658649

**Authors:** Mythili Subharam, Ryan Koehler, Tejas Sreedhar

## Abstract

**PURPOSE:** Early detection of hepatocellular carcinoma (HCC) is critical for improving patient outcomes, yet current screening tools lack sensitivity and specificity. We demonstrate a flexible machine learning framework for HCC detection using methylation profiles from bisulfite sequencing across multiple assay platforms and sample types. The framework supports a “split-and-filter” approach that routes each sequenced sample to an assay-matched classifier without requiring cross-assay feature compatibility.

**PATIENTS AND METHODS:** We constructed assay-specific classifiers using four independent public methylation datasets (∼2,500 total samples) representing distinct bisulfite-sequencing technologies: GSE93203 (MCB-targeted hypermethylation), GSE63775 (MCTA-Seq tandem-repeat hypermethylation), PRJCA001372 (HBV-integration–associated hypomethylation), and the HCC subset of GSE149438 (EpiPanGIDX - DMR-level hyper- and hypomethylation). Separate models were trained using using the biologically relevant features for each dataset and evaluated in two independent blind validation datasets: published tissue WGBS (24 samples: 12 early-stage HCC, 12 matched controls; PRJNA984754) and a new preclinical plasma cfDNA WGBS dataset (12 samples) generated at a commercial sequencing laboratory.

Limited feature overlap among assays precluded a single unified model. Instead, overlapping features enabled construction of a proof-of-concept meta-classifier for sample routing across assay-specific models.

**RESULTS:** Assay-specific cfDNA models, trained independently on CpG sites from original publications, were evaluated using the biopsy(tissue) dataset and a new plasma dataset as blind validation. All four assay-specific models generalized well to the validation data, with accuracies of 83.5%-100%. In the validation with the plasma cfDNA samples, the best-performing classifier (among XGBoost, Random Forest, and Logistic Regression) for each public dataset achieved 80–100% sensitivity and 86–100% specificity, with all Stage 2 cases correctly detected across models. The single Stage 1A case showed methylation levels overlapping with cirrhotic controls, consistent with biological expectations. Despite this, a couple of the models predicted this correctly, showing greater sensitivity to Stage 1 cancer.

**CONCLUSION:** A generalizable framework for early detection of HCC composed of assay-specific classifiers and a meta-classifier is described. This architecture readily accommodates addition of new assays via feature-matched models and meta-classifications. Larger, prospectively collected studies are necessary to confirm performance and enable clinical translation.

## INTRODUCTION

Hepatocellular carcinoma (HCC) is the most common form of primary liver cancer and a leading cause of cancer-related mortality worldwide. Prognosis is closely tied to stage at diagnosis, yet the majority of cases are identified at advanced stages, when curative treatment options are limited. The current standard of care for early HCC surveillance in high-risk patients is ultrasound imaging with or without serum alpha-fetoprotein (AFP) testing, but this offers only modest sensitivity (∼65%), particularly for early-stage tumors^7^ and obese patients^8^. There is a pressing need for more sensitive, non-invasive detection methods that can identify HCC at an earlier, curable stage.

Liquid biopsy approaches based on cell-free DNA (cfDNA) have shown considerable promise for non-invasive cancer detection. While many commercial platforms^25^ focus on detecting somatic mutations, especially for MRD/recurrence, this strategy is poorly suited for early detection in HCC, where tumor-derived DNA often constitutes a small fraction of total cfDNA. Moreover, mutational profiles vary widely across patients and disease stages, making it difficult to train models that capture consistent, generalizable patterns^9,10^. In contrast, epigenetic DNA methylation level alterations are early events in tumorigenesis and tend to occur in conserved regulatory regions across individuals.

These patterns are more stable across patients and cancer stages, providing a more consistent signal for machine learning–based classification^11^. Bisulfite sequencing of cfDNA enables single-base resolution of methylation patterns and can be applied in both genome-wide and variously targeted assay formats. Compared to array-based or probe-constrained methods, bisulfite sequencing offers greater sensitivity and flexibility in detecting tumor and tissue-specific epigenetic changes, even in samples with low tumor DNA content, which makes it particularly well suited for early cancer detection^11^.

Other methylation assay formats such as MeDIP-seq offer cost advantages compared to bisulfite sequencing. However, non-specific antibody binding to common sequence motifs, such as short tandem repeats, can cause a high rate of false positives in MeDIP-seq^24^, a problem not present in bisulfite sequencing. Several recent studies have advanced cfDNA-based detection of HCC using alternative epigenomic signals.

Fragmentation-based approaches leverage cfDNA fragment end-motifs^20^ or genome-wide fragmentomes to detect cancer-associated nucleosome patterns^21^. Other studies have demonstrated methylation signatures and multi-omic mutation/methylation assays with promising accuracy. However, these methods generally rely on specialized sequencing constructs, proprietary fragmentation pipelines, or fixed targeted panels that are not directly interoperable across heterogeneous assay types. In contrast, our goal in this work is not to introduce a new cfDNA biomarker modality but to establish a generalizable, assay-flexible machine learning framework capable of handling diverse bisulfite-sequencing datasets. This architecture allows incorporation of heterogeneous markers, including those from future studies, while maintaining compatibility with genome-wide WGBS data used in early-stage clinical development.

While cfDNA methylation is a promising biomarker class for early cancer detection^1^, published models and commercial tests have struggled to generalize beyond their original datasets. For example, the Galleri methylation-based multi-cancer detection test demonstrated high specificity (∼99%) but only 16.8% sensitivity in early-stage cancers^15^. We hypothesized that assay-aware machine learning models, trained on biologically grounded methylation features, could achieve high accuracy across different bisulfite sequencing platforms, sample types and patient populations. Use of an ensemble approach, specifically a "mixture of experts" or a "gating network" architecture, might yield a sensitive and general predictive tool^12^.To test this, we developed a modular framework using public cfDNA datasets generated with four distinct bisulfite sequencing assays. We then evaluated generalizability via blind validation against an independent public tissue-based dataset, as well as a newly tested small cfDNA plasma biosample cohort.

## METHODS

### Data Sources

We analyzed four publicly available bisulfite sequencing datasets representing plasma-derived cfDNA from hepatocellular carcinoma (HCC) patients and healthy controls, each generated using a distinct assay (**Table 1**). These include: GSE93203, which used targeted bisulfite sequencing of plasma-derived cfDNA targeting methylation-correlated blocks (MCBs) across 467 short genomic regions^4^. These regions were selected based on coordinated methylation patterns associated with HCC, but this small footprint may be poorly sampled by broader sequencing assays; GSE63775, another targeted assay with limited genomic footprint, which used Methylated CpG Tandem Amplification and Sequencing (MCTA-Seq) of plasma-derived cfDNA to enrich for fully methylated CpG islands^2^; PRJCA001372, which used low-depth whole-genome bisulfite sequencing (WGBS) of plasma-derived cfDNA targeting hypomethylated markers^3^; and EpiPanGIDX^19^, a targeted methylation panel covering multi-GI cancer signatures from which only HCC samples + controls were used. Features included HCC-specific hyper- and hypomethylated DMRs (differentially methylated regions).

**Table 1.** Model summary.

| Model ID | Assay Type | Selected Features | Number of Samples | K-fold Internal Validation Accuracy | Blind Validation (PRJNA984754) | AUC | Sensitivity | Specificity |
| --- | --- | --- | --- | --- | --- | --- | --- | --- |
| GSE63775 | Targeted MCTA-Seq (cfDNA) | 3,988 CpG sites | 151 | 84.1% (avg. 5-fold) | 83.3% | 0.80 | 83.3% | 83.3% |
| GSE93203 | MCB Targeted Bisulfite (cfDNA) | 32 CpG sites | 2191 | 93.5% (avg. 5-fold) | 83.3% | 0.875 | 91.7% | 75% |
| PRJCA001372 | Low depth WGBS (cfDNA) | 3,592 CpG sites | 60 | 88.9% (avg. 5-fold) | 100.0% (threshold=0.8) | 1.00 | 100.0% | 100.0% |
| GSE149438 (EpiPanDX) | Targeted hypo and hypermethylated DMRs | 435 DMRs (differentially methylated regions) | 98 | 90% (avg, 5 fold) | 83.3% | 0.80 | 83.3% | 83.3% |
| PRJNA984754 | WGBS<br>(Biopsy<br>tissue) | Three<br>variants:<br>union<br>(2,000),<br>hypo-only,<br>hyper-only | 24 | 100.0% (3-fold,<br>union@0.7<br>threshold) | — | — | — | — |

1. All models used XGBoost classifiers, which outperformed Random Forest in internal comparisons. 2. “Selected Features” indicates the number of CpG sites retained after filtering and preprocessing. 3. For PRJNA984754, three models were trained using different feature sets: **Hypo-only:** CpG sites selected based on hypomethylation in PRJCA001372 **Hyper-only:** CpG sites selected based on hypermethylation in GSE63775 **Union:** Top 2,000 combined features from both hypo- and hypermethylated site sets

Two datasets were used for independent validation of each of the trained models. PRJNA984754, a moderate-depth WGBS dataset of matched HCC and adjacent normal liver tissue biopsy samples^5^ and preclinical validation with 12 plasma samples (5 early stage HCC, 7 controls (5 CLD; 2 healthy), sequenced using Illumina NovaSeq X, targeting 200M paired-end reads, 150 bp, performed at Zymo Research.

### Feature Selection

For each dataset, FASTQ sequencing files were processed through a bioinformatics pipeline using Bismark^16^ to compute “beta values”, defined as the ratio of methylated reads to total reads covering a CpG site. All sites with aligned reads were included in beta value calculations. Initial data matrices were created by collecting beta values from all samples at CpG sites previously identified as relevant for HCC detection by the original study authors. These were subsequently filtered to remove CpG sites exhibiting zero variance across samples.

The GSE93203 (MCB) dataset yielded limited CpG coverage when raw FASTQ files were processed through our standard pipeline, with many loci from the original study showing little or no usable signal. We then obtained the original study’s published beta value matrix (n=467 CpGs) and manually mapped the biomarker coordinates described in the paper to genomic locations. These mapped regions were cross-referenced against the 467 CpGs in the matrix, resulting in a final set of 32 CpG sites corresponding to 18 reported biomarker loci.

### Model Training and Internal Validation

Because each cfDNA dataset was generated from a different assay, overlap in CpG coverage across the datasets is poor. This precluded construction of a robust unified model using only shared features. Instead we opted to train four separate machine learning models, using CpG sites selected from the corresponding publications. While the reduced feature set across all models was not adequate for joint modeling, it nevertheless proved sufficient to support a meta-classifier for model routing of samples to appropriate assay-specific models.

The XGBoost^18^ algorithm is known for its strong performance, especially when handling missing values without requiring imputation^13^. We developed models specific for each explored dataset using this approach. Random Forest classifiers and Logistic Regression classifiers were also evaluated but did not match XGBoost performance in most cases. Internal validation was performed using k-fold cross-validation (K = 5 or 10, based on dataset size), with a default classification threshold of 0.5 unless otherwise noted.

### Validation

#### a) Blind Validation with tissue dataset

To assess generalizability, each cfDNA model was applied to the independent PRJNA984754 tissue biopsy dataset. For each model, features were filtered to match the selected features of the respective training model. All models were evaluated using k-fold cross-validation and standard performance metrics (accuracy, sensitivity, specificity, and AUC).

Predictions were generated using each trained classifier. For each model, a default classification threshold of 0.5 was initially applied. Where appropriate, thresholds were adjusted to account for the beta value range shifts between the training and test datasets, due to assay differences.

### Hybrid Tissue Model Construction

To explore the predictive value of features identified from cfDNA in a biopsy tissue context, another model was trained directly on PRJNA984754. This model used CpG sites selected from the PRJCA001372 (hypomethylated, low depth WGBS) and GSE63775 (hypermethylated, MCTA-Seq) datasets. Three variant subsets were assessed: hypomethylated-only features, hypermethylated-only features, and a combined hybrid model using the top 2,000 sites after filtering. To select the top 2,000 features, an initial XGBoost model was trained on the full filtered feature set, and feature importance scores were extracted. The 2,000 highest-ranking features were then used to train a second, reduced model for performance evaluation. This model’s performance was assessed using 3-fold cross-validation.

#### b) Preclinical validation for the Plasma sample dataset

##### Wet-lab preprocessing

The 12 plasma samples underwent: **cfDNA extraction:** Zymo Quick-cfDNA Serum & Plasma Kit; **Bisulfite conversion:** EZ DNA Methylation-Lightning Kit (Zymo Research); **Library preparation:** Zymo-Seq WGBS Library Prep Kit; **Sequencing platform:** Illumina NovaSeq X **Sequencing depth:** Target 200M paired-end reads (150 bp); **Alignment Genome:** HG19 Fastq and BAM files were delivered to the OmicXHealth preprocessing pipeline where they were converted to cov.gz files to build the methylation beta value matrices with samples as rows and features as columns for each model. Each WGBS validation sample was filtered to only the selected features/regions of each model before classification.

##### Normalization methods for batch correction

To account for assay differences between the training and validation datasets, the following normalization methods were applied to shift the validation dataset values to the range of the training dataset. No changes to the training datasets were made.

Depending on assay: for MCB (GSE93203): Mean-shift alignment only; β-values used directly (M-values provided no improvement) for **MCTA-Seq (GSE63775):** β → M transformation; Reference-anchored ComBat (training as reference) **HBV-hypomethylation models (PRJCA001372):** Region-level model: median-shift alignment; CpG-discrete model: no normalization; NaNs preserved **EpiPanGIDX (GSE149438):** Mean-shift alignment to training distribution. Thresholds via 5-fold median Youden-J.

##### Model training

For each dataset, we trained: **XGBoost, Random Forest, Logistic Regression (with class weights).** Table 2 shows the validation performance of each model.

**Table 2 -.**
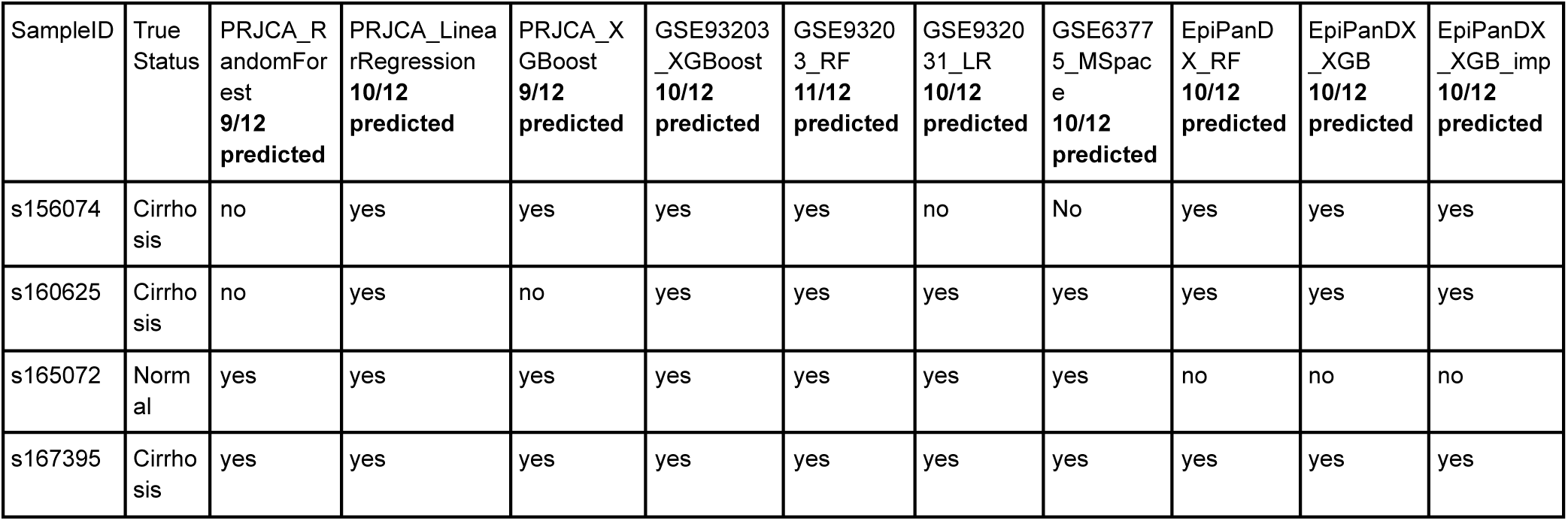
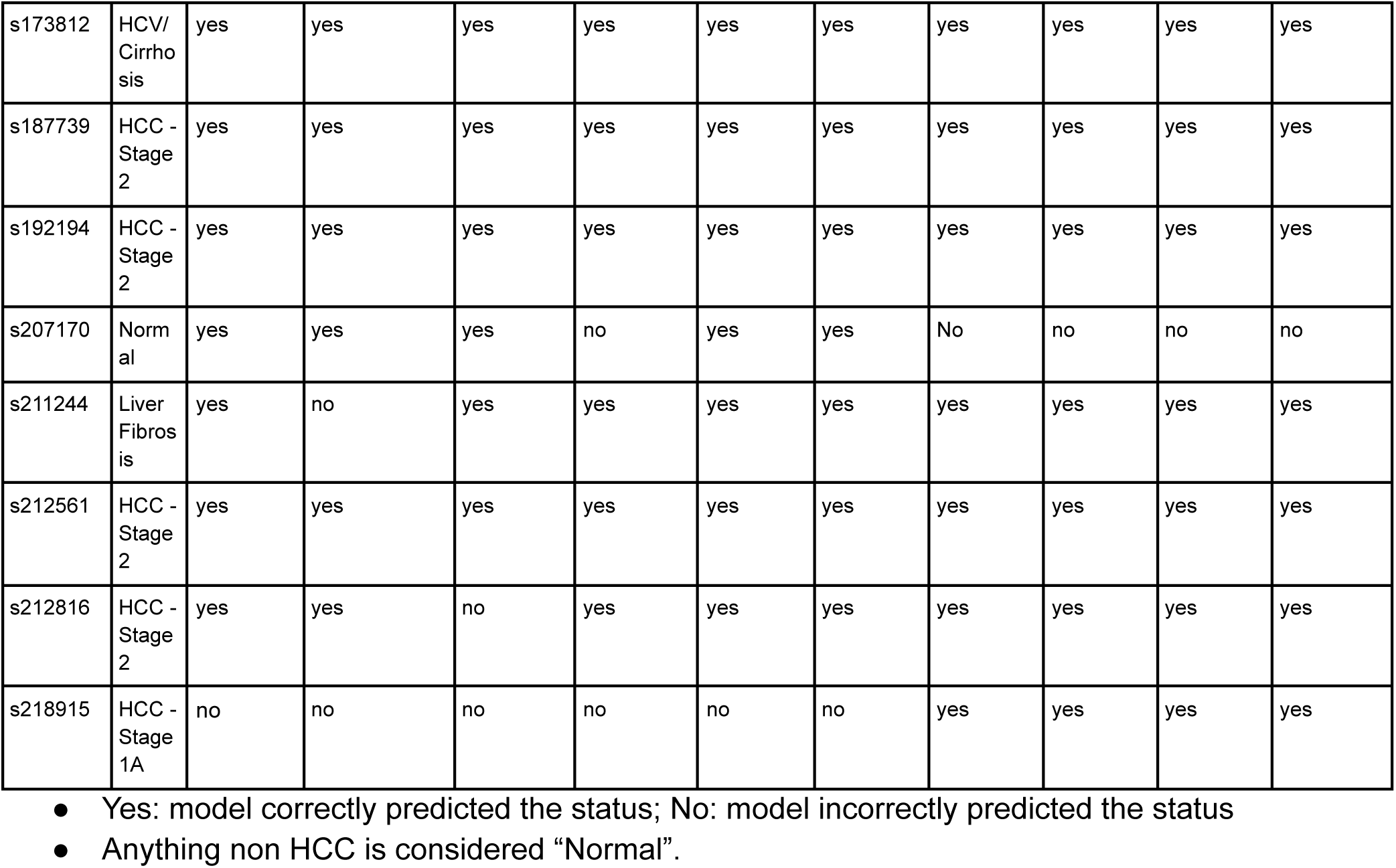
Results of Pre-clinical validation with biosamples - How each Model predicted each sample.

## RESULTS

All models achieved strong internal performance, with mean k-fold cross-validation accuracy of 84%-94%. To assess generalizability, we performed blind validation using an independent tissue biopsy WGBS dataset consisting of 12 matched liver cancer-normal tissue pairs (24 samples total). Model inputs and performance metrics are summarized in **Table 1** (and ROC curves are included as supplementary figures).

### Key Findings for the pre-clinical validation with plasma samples (high-level)

- ● Using the **best-performing classifier for each dataset**, achieved:

○ **Sensitivity:** 80–100%
○ **Specificity:** 86–100%
- ● **All Stage 2 HCC samples** were detected across all models
- ● The **Stage 1A HCC** sample consistently overlapped with cirrhotic controls across methylation markers (expected in cfDNA biology)
- ● False positives occurred **only in healthy normals**, which are **not part of the intended-use population** (the clinical population = CLD/HBV/HCV high-risk cohort)

### Model level performance

**GSE63775 (Promoter hypermethylation model)**

● **Sensitivity: 100%**
● **Specificity: 71%**
● Best overall separability after ComBat-based normalization
● Directionality consistent in **all** Zymo HCC samples

**GSE93203 (32 CpG MCB model)**

● **Sensitivity: 80%**
● **Specificity: 100% (RF classifier)**
● Stability improved via retrospective Youden-J calibration

**PRJCA001372 (HBV hypomethylation models)**

● **Sensitivity: 60–75% depending on region-level vs CpG-level**
● **Specificity: 86%**
● Strongest detection in deeper hypomethylated Stage 2 cases

**EpiPanGIDX HCC DMR model**

● **Sensitivity: 100%**
● **Specificity: 71%**
● Robust cross-platform consistency despite assay differences

### Blind validation performance of PRJNA984754 dataset

**GSE63775 (MCTA-Seq)**

**Threshold Used:** 0.45 *(see Key Insights for rationale)*

**Accuracy on Blind Validation:** **83.3%**

**Key Insights:** The CpG site–level model trained on GSE63775 generalized well to the independent PRJNA984754 dataset, achieving high accuracy despite differences in assay type (targeted vs. WGBS) and sample source (cfDNA vs. tissue). While global methylation distributions show broader overlap between cancerous and healthy samples in the validation set (**Figure 1B**), site-level boxplots (**Figure 2A**) reveal strong discriminative power at key CpGs. Clear hypermethylation trends in cancer are preserved across datasets, suggesting that the model’s predictive performance is driven by a focused subset of informative CpG sites rather than global patterns.

**Figure 1.**
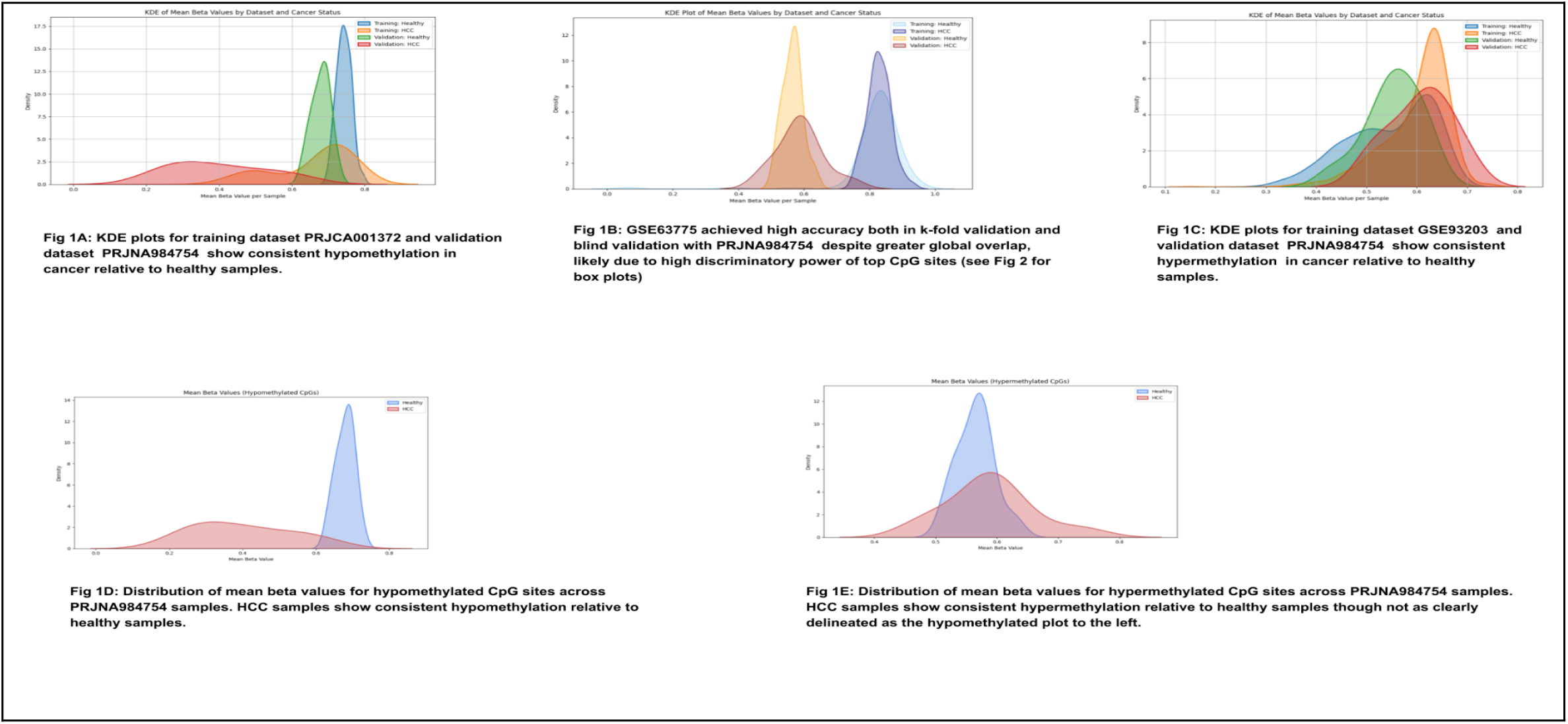
KDE Distribution Plots for Global and Site-Level Methylation Trends. Kernel density estimates (KDEs) comparing the distribution of beta values between cancerous and healthy samples.

**Figure 2.**
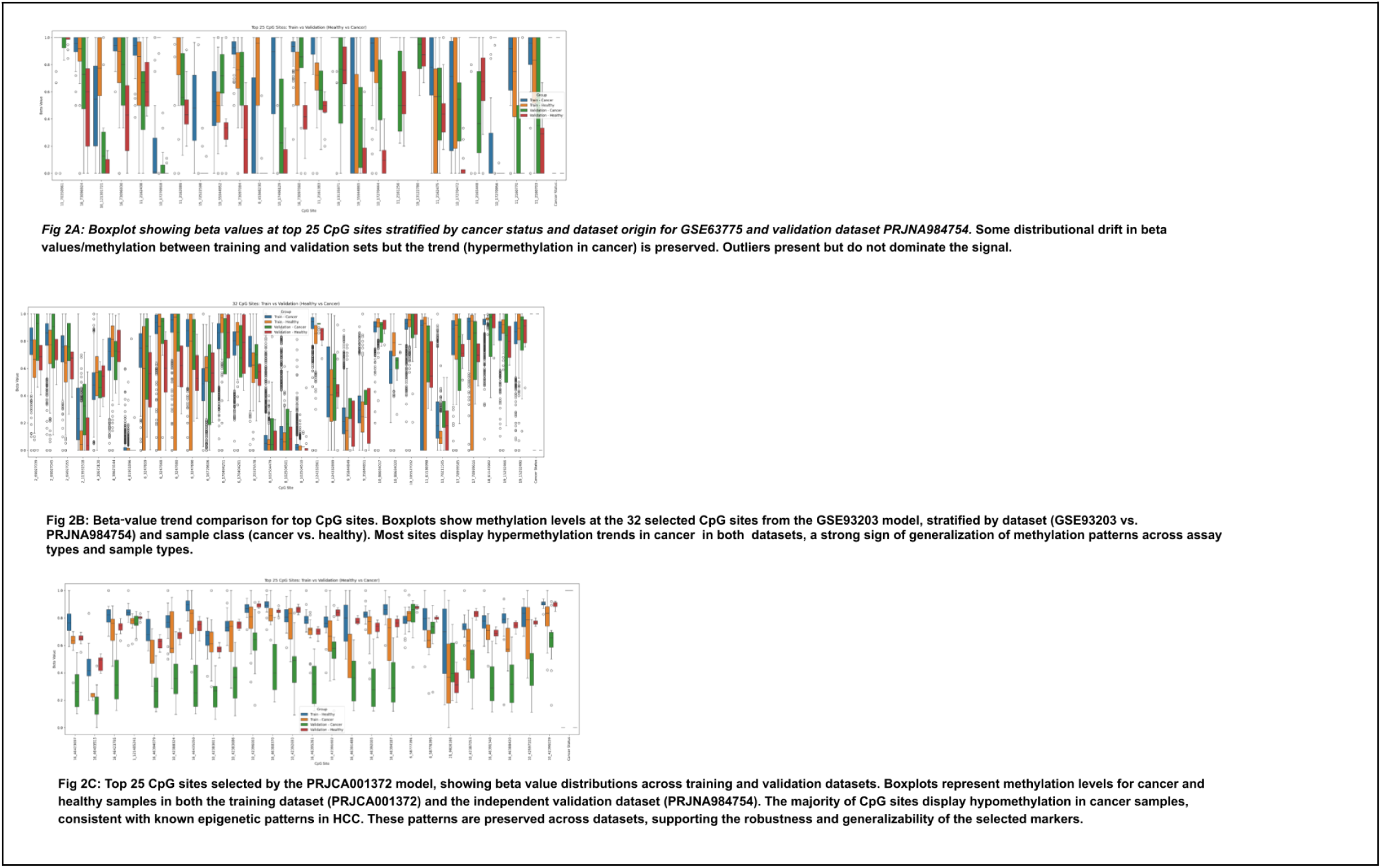
Boxplot Summary of Methylation Differences Across Training and Validation datasets. Boxplots comparing beta values of cancerous and healthy samples at key CpG sites across training and validation datasets.

Beta values in the validation dataset are shifted lower across the board—likely reflecting biological or technical differences between plasma-derived and tissue-derived samples. This required a lower prediction threshold to retain sensitivity. The model, having learned “high beta = cancer,” needed a threshold adjustment to correctly identify cancer samples with comparatively lower beta values in the validation set.

**GSE93203 (MCB-Based Targeted Sequencing)**

**Threshold Used:** 0.5 (after batch correction)

**Accuracy on Blind Validation:** 83.5%

**Key Insights:**

The GSE93203 model, trained using a refined set of 32 CpG sites mapped from 18 published biomarkers, achieved strong generalization to the independent PRJNA984754 validation dataset following status-aware batch correction. While earlier attempts using larger feature sets (e.g., 467 CpGs from the supplementary matrix) and traditional batch correction approaches failed to generalize, restricting the model to biologically validated markers improved both stability and performance.

After applying a cancer status–specific mean shift correction to the methylation beta values—where mean differences between cancerous and non-cancerous samples in the training and validation sets were aligned separately—a clear separation between classes emerged as seen in **Figure 1C**. Fixing the classification threshold at 0.5 post-correction yielded balanced performance across cancerous and non-cancerous tissue samples. This result suggests that although the original assay used targeted enrichment of MCB regions, a focused subset of biologically validated CpG markers can generalize effectively across platforms when normalized appropriately.

**PRJCA001372 (Low-Depth WGBS)**

**Threshold Used:** 0.8 (see Discussion for rationale)

Accuracy on Blind Validation: **100%**

**Key Insights:**

This model, trained on cfDNA-derived hypomethylated CpGs, achieved perfect classification when tested against PRJNA984754 tissue samples. KDE plots (**Figure 1A**) confirm that cancer samples have lower global beta values, consistent with expected hypomethylation in HCC. The site-level boxplots (**Figure 2C**) reinforce this, showing preserved hypomethylation across training and validation sets.

Although overall methylation levels in the validation dataset were lower for both healthy and cancer samples, cancer-specific hypomethylation remained distinct. A higher threshold was required to correctly separate the classes, consistent with the model’s learning that “low beta = cancer.” This underscores the robustness of hypomethylation-based models across different sequencing platforms and sample types.

**Standalone Models Trained on PRJNA984754 dataset (WGBS tissue dataset) Accuracy (3-Fold Cross-Validation):**

- ● Hypomethylated-only features (from PRJCA001372): 23/24 correct (threshold
- = 0.5)
- ● Hypermethylated-only features (from GSE63775): 20/24 correct (threshold = 0.5)
- ● Hybrid (hypomethylation + hypermethylation features): 100% (threshold = 0.7)

Key Insights:

To test whether externally derived feature sets retain predictive power in a purely tissue-derived context, we trained three PRJNA984754-only models using CpG sites selected from the other datasets. KDE plots (**Figure 1D and E**) show meaningful class separation for both hypo- and hypermethylated features. Panel 1D (hypo) reveals clean separation between HCC and healthy samples, while Panel 1E (hyper) shows greater overlap—though cancer samples remain hypermethylated (skewed toward higher beta values).

Boxplots for hypermethylated CpGs (**Figure 2A**) indicate that individual sites maintain strong trends even when global distributions are less distinct. These findings suggest that while global hypomethylation offers broader signal, select hypermethylated markers can offer high site-level resolution. The hybrid model combining both achieved perfect classification, reinforcing the value of integrating feature types when sequencing depth allows.

ROC curve analysis (see Supplementary section) further illustrates the generalizability of models across datasets. The GSE63775-based model achieved an AUC of 0.9059 on blind validation samples, misclassifying only 4 out of 24. The GSE93203 model yielded an AUC of 0.875 misclassifying 4 out of 24.Notably, the PRJCA001372 model, trained and validated on WGBS-based data, achieved an AUC of 1.0, suggesting near-perfect alignment and underscoring the impact of methodological consistency between training and validation.

### Overall Validation Summary

All cfDNA-based models — PRJCA001372 (low-depth WGBS), GSE63775 (MCTA-Seq), GSE93203 (MCB-based targeted bisulfite sequencing) and EpiPanGIDX (targeted DMRs)— demonstrated strong generalizability to the independent PRJNA984754 tissue WGBS validation set(24 samples) and to the Zymo plasma cfDNA preclinical validation dataset(12 samples) . Although trained on plasma-derived cfDNA data using different sequencing methods and distinct feature sets, each model successfully distinguished cancerous from non-cancerous tissue samples after aligning methylation profiles across datasets. This supports the hypothesis that biologically relevant methylation patterns at carefully selected individual CpG sites are robust to both biospecimen type and sequencing platform.

The GSE93203 model initially struggled during validation. Despite high intra-dataset accuracy (∼94%), it misclassified all healthy samples as cancerous when applied directly to the validation set. Ultimately, a refined model restricted to 32 CpG sites that mapped to the originally published markers was constructed. A cancer status–specific mean shift correction was applied to the beta values, achieving stable separation between classes and a prediction accuracy of 83.3%. This result suggests that when focused on a validated subset of biomarkers and normalized appropriately, even assay-specific models can generalize well across platforms.

Prediction thresholds required for accurate classification varied across models due to global shifts in methylation levels in the tissue-derived validation set. As shown in **Figure 2** (Panels A–C), the directionality of cancer-associated methylation shifts remained consistent even when absolute beta values differed, facilitating successful generalization across the three models.

### Cross-Platform Signal Reproducibility

Across four unrelated public datasets spanning four assay chemistries (MCB, MCTA-Seq, WGBS, targeted-panel), the underlying HCC methylation patterns—hypermethylation in cancer-associated regions or hypomethylation near HBV integration loci—were preserved during validation on two independent datasets.

The fact that tissue WGBS (PRJNA984754), and cfDNA WGBS from a commercial lab (Zymo) **both reproduced the expected biological directionality** validates the markers though larger studies are required to confirm generalizability.

### 3.2 Preclinical Feasibility Study Interpretation

The cfDNA validation confirms **Hyper**methylation markers (GSE93203, GSE63775, EpiPanGIDX) generalize more strongly than hypomethylation markers. Stage 2 HCC is readily detectable Stage 1A presents some overlap with cirrhotic controls. One model had false positives only in **healthy** normals, potentially because it hadn’t been trained on any truly healthy samples,which **are not the intended clinical cohort.** This supports the interpretation that assay differences and low tumor fraction, not model inadequacy, drive the remaining misclassifications.

While overall classification accuracy (83–92%) reflects meaningful separability, performance is still limited by **batch-dependent shifts** between training and validation datasets.

The isotonic-calibrated models required minor threshold recalibration (via Youden-J optimization) to achieve high sensitivity and specificity, suggesting that the discriminative biology is present but the probabilistic calibration remains assay-specific.

This reinforces the view that the primary limitation at this stage is assay normalization, not model structure or biomarker quality. If the calibrated threshold values from this batch hold in the next batch, then these public dataset trained models themselves could be used for CLIA-grade assay validation for commercialization.

## DISCUSSION

This study demonstrates that a generalizable machine learning (ML) framework can accurately detect hepatocellular carcinoma (HCC) using cfDNA methylation data from multiple sequencing platforms. Unlike prior studies that relied on siloed datasets or proprietary assays, our approach integrates platform-specific models within a unified meta-classifier architecture. Each model was trained on biologically relevant CpG markers identified by the original study authors, using conservative feature filtering to balance granularity and generalization. Cross-validation showed strong within-cohort performance, and blind validation using an independent whole-genome bisulfite sequencing (WGBS) liver tissue dataset confirmed that cfDNA methylation signatures are both biologically robust and technically generalizable.

A key feature of this framework is its flexibility in handling assay heterogeneity. When the sequencing method is known (e.g., WGBS), sample data can be preprocessed into multiple filtered versions—each corresponding to the feature set of a trained model—and passed through all models independently. Final classification is based on the prediction with the highest confidence score, which reflects how well the sample’s methylation pattern matches disease-associated features. For example, strong hypomethylation across key sites yields higher confidence from the hypomethylation model. In real-world scenarios where assay metadata may be unavailable, the meta-classifier offers a robust alternative: using a small, shared subset of intersecting CpG features, it routes the sample to the most appropriate model based on methylation signature similarity—enabling accurate predictions without prior knowledge of sequencing modality.

This work also confirms the stability of methylation patterns across cancer stages and sample types. All four training datasets (MCTA-Seq, WGBS, targeted DMR, and padlock probe–based assays) included early to late HCC stages, yet each model performed well during cross-validation. The WGBS liver tissue dataset, despite being derived from solid tissue rather than plasma, validated strongly against all three cfDNA-trained models, suggesting that cfDNA and tissue share stable methylation signatures across sequencing methods and clinical contexts.

Another insight was the impact of sequencing technology on model generalizability. Among the three models, the one trained on WGBS achieved the highest accuracy during blind validation, likely reflecting the benefit of matched sequencing platforms between training and test sets. Still, the MCTA-Seq and padlock-based models also performed well despite their differing platforms, underscoring that methylation-based classifiers can generalize across assays when features are biologically meaningful.

We also observed that methylation beta values often shift downward in absolute magnitude between training and validation datasets, likely due to differences in sample type (plasma vs. tissue) and sequencing depth. For the PRJCA001372-trained model, this global downward shift caused all validation samples to appear more hypomethylated than expected. While cancer samples still had lower methylation than controls, all probabilities shifted upward. Raising the classification threshold restored accuracy by separating the classes based on the new distribution. The GSE93203-trained model showed a more extreme mismatch: both cancerous and healthy validation samples fell into the non-cancer methylation range of the training data. Although relative trends were preserved (cancer < healthy), the shift invalidated threshold-based classification. In this case, batch correction was necessary to realign distributions and recover predictive accuracy. These findings suggest that while modest platform shifts can be addressed via threshold tuning, substantial mismatches require explicit harmonization.

## CONCLUSION

This study demonstrates the feasibility of a generalizable machine learning framework for early hepatocellular carcinoma (HCC) detection using cfDNA methylation data derived from diverse bisulfite sequencing platforms. We trained three distinct models using public datasets capturing hypermethylation, hypomethylation, and hybrid signatures, and validated their performance on an independent WGBS liver tissue dataset. All models achieved strong predictive accuracy, with the hypomethylation model reaching 100% accuracy after threshold adjustment.

Because all models operate on CpG-level features, the framework allows flexible routing of individual samples through multiple classifiers based on their methylation profile. This architecture enables scalable and cost-effective deployment across a range of sequencing protocols.

Crucially, the framework is extensible. As emerging cfDNA methylation biomarkers are identified in the literature—such as GJA4 in HBV-associated HCC—they can be incorporated into future iterations of the model without requiring changes to the assay protocol. This adaptability strengthens long-term clinical utility.

Collectively, these findings support the use of a multi-model, meta-classifier–based approach for HCC detection, offering both technical flexibility and biological robustness. Retrospective pre-clinical validation using biobank-derived cfDNA samples assessed real-world performance but larger studies are required to inform potential integration into early cancer screening workflows.

## Acknowledgments

We thank Paul Gaudin and Dr. Elena Tomas Bort for their early contributions to the data preprocessing and feasibility validation phases of this project.

## Funding Information/Research Support

No external funding was received. The research funding was bootstrapped by the corresponding author on behalf of OmicXHealth.

## Disclaimers

None.

