## Supplementary tables and figures for "A Generalizable Machine Learning Framework for cfDNA based Early Detection of Hepatocellular Carcinoma: a Feasibility Study with Preclinical Validation"

### Supplementary Section

### Tables

**Table S1.** Public datasets used in study for model training and validation.

| Dataset Accession | Publication | Tissue | Assay | Samples | Biomarkers / features in original study | Sequence data features |
|---|---|---|---|---|---|---|
| GSE63775 | Wen 2015[2] [Wen2015] | cfDNA | MCTA-Seq | 151 total; 91 plasma: (36 HCC; 16 cirrhosis; 39 normal) Tissue: 60 (27 HCC + adjacent tissue, 3 normal liver) | Tumor-specific hypermethylation CpG islands in plasma and tissues; identified 382 differentially methylated CGIs (pdmCGIs) for early-stage detection. | Cell free DNA. Tumor-specific hypermethylation CpG islands in plasma and tissues. Assay uses multi-3'CG primers. Identified 382 pdmCGIs for early-stage detection. "Methylated CpG tandems amplification sequencing" (MCTA-seq) |
| GSE93203 | Xu 2017[4] [Xu2017] | cfDNA | MCB Targeted Bisulfite seq | 2,191 total; 1,221 HCC; 970 normal | 10 MCBs (Methylation Correlated Blocks) | Cell free DNA. Targeted Bisulfite Sequencing "Methylation Correlated Blocks" (MCBs). Assay uses padlock probes. |
| PRJCA001372 | Zhang 2020[3] [Zhang2020] | cfDNA | Low depth WGBS | 60 total; 26 HCC; 34 non-HCC but chronic liver disease (Hepatitis / Cirrhosis) | 28 hypomethylated regions associated with Hepatitis B Virus (HBV) integration, selected from 5,851 initial sites based on methylation difference, sequencing depth, and | Cell free DNA. Low coverage whole genome bisulfite sequencing. HBV integration-associated hypomethylation regions. |

| | | | | | | |
|---|---|---|---|---|---|---|
| | | | | | correlation with tumor tissue. | |
| PRJNA984754 | Zhao 2023[5] [Zhao2023] | Tissue biopsy | WGBS | 24 samples, 12 HCC and 12 healthy | None | Matched tumor and normal adjacent tissue pairs. |

### AUC-ROC curves

**Supplementary Figure S1. ROC Curve for XGBoost model trained on GSE93203 (Hypermethylated Sites)**

Blind validation was performed on PRJNA984754. Its validation AUC was **0.875** suggesting good external generalization.

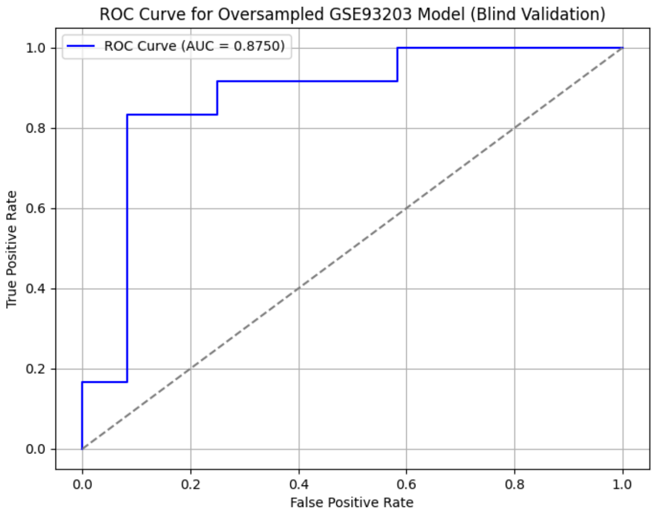


**Supplementary Figure S2. ROC Curve for XGBoost model trained on GSE63775.**

Blind validation was performed on PRJNA984754. The model was trained on individual

CpG sites mapped from MCTA-Seq derived regions. The AUC was **0.9059**, showing strong predictive performance and generalization to tissue-derived WGBS samples.

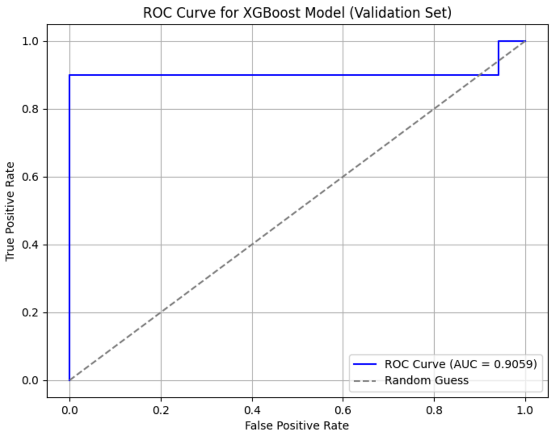


**Supplementary Figure S3. ROC Curve for PRJCA001372-trained model on PRJNA984754 blind validation samples.**

The model trained on cfDNA samples using hypomethylated CpGs achieved **AUC = 1.000**, demonstrating perfect discrimination on this validation dataset. However, the sharp performance suggests potential data alignment with PRJNA984754 methylation patterns or shared biological signal.

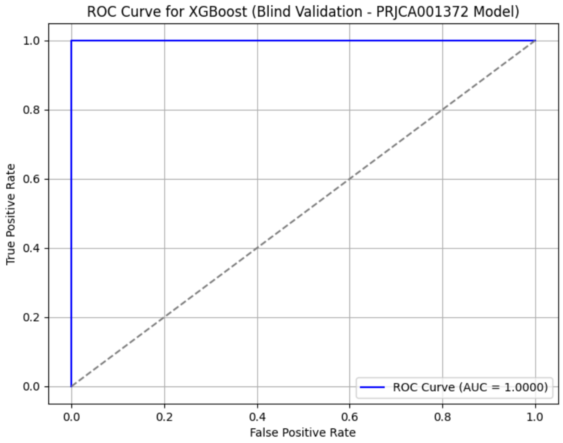


**Supplementary Figure S4. ROC Curve for PRJNA984754 self-model trained using hypomethylated features from PRJCA001372.**

The AUC was **0.9271**, indicating that these hypomethylated markers translated well from cfDNA to tissue WGBS in the same disease context.

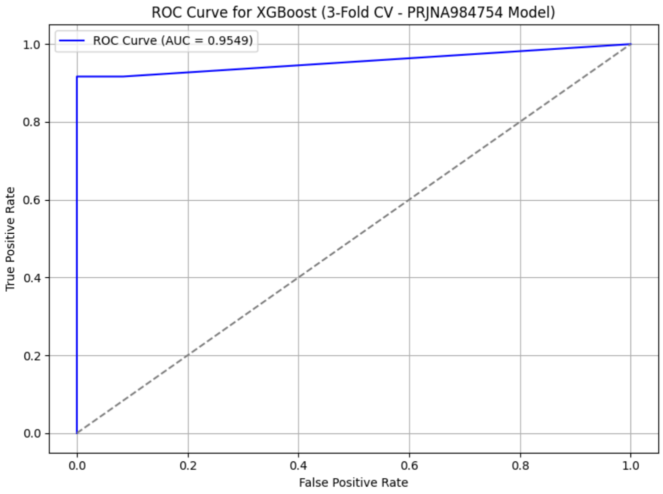

**Supplementary Figure S5. ROC Curve for PRJNA984754 self-model trained using hypermethylated features from GSE63775.** This model was trained and evaluated using 3-fold cross-validation on PRJNA984754 tissue samples, with features selected from hypermethylated CpG sites originally identified in GSE63775. Although the mean classification accuracy across folds was 83.3%, the ROC-AUC was 0.9549, indicating strong rank-order separation between HCC and healthy samples. This discrepancy highlights how models with modest classification thresholds can still achieve high discriminative performance.

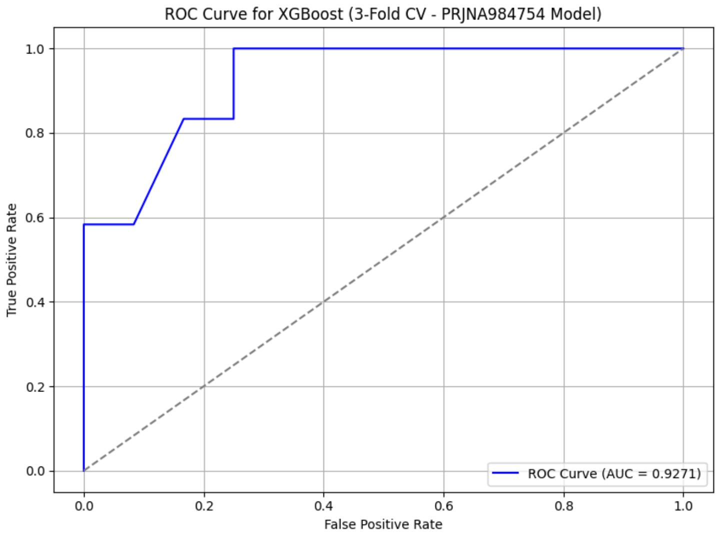


**Supplementary Figure S6. ROC Curve for PRJNA984754 self-model trained using union of top 3000 hyper + hypo CpG features.**

AUC = **1.000**, confirming strong internal validation when combining diverse methylation markers from both GSE63775 and PRJCA001372.

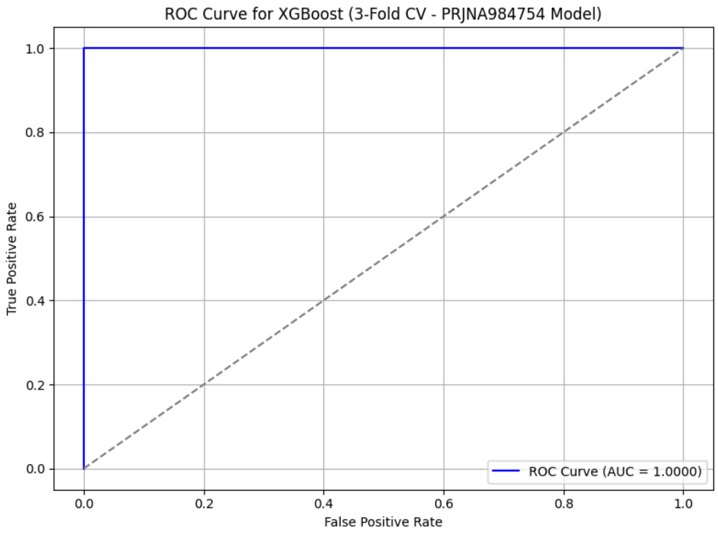
ROC Curve for XGBoost (3-Fold CV - PRJNA984754 Model)
True Positive Rate
False Positive Rate
0.0
0.2
0.4
0.6
0.8
1.0
ROC Curve (AUC = 1.0000)
